# Structure of the F-tractin–F-actin complex

**DOI:** 10.1101/2024.10.07.616643

**Authors:** Dmitry Shatskiy, Athul Sivan, Roland Wedlich-Söldner, Alexander Belyy

**Affiliations:** Membrane Enzymology Group, Groningen Institute of Biomolecular Sciences and Biotechnology (GBB), Faculty of Science and Engineering, University of Groningen, Groningen, The Netherlands; Institute of Cell Dynamics and Imaging, and Cells-in-Motion Interfaculty Center (CiMIC), University of Münster, Münster, Germany

**Author notes:** The authors contributed equally (Dmitry Shatskiy, Athul Sivan).

## Abstract

F-tractin is a short peptide widely used to visualize the actin cytoskeleton in live eukaryotic cells. Similar to other actin-binding probes, F-tractin alters actin organization and impairs cell migration when expressed at high levels. In addition, the probe has been reported to directly induce actin bundling. To elucidate the mechanism behind these effects, we determined the structure of the F-tractin–F-actin complex using electron cryo-microscopy. Our analysis revealed that the F-tractin peptide consists of a flexible N-terminal region and an amphipathic C-terminal helix. The N-terminal part is completely dispensable for F-actin binding but is responsible for the actin bundling effect. The C-terminal helical region interacts with a hydrophobic pocket formed by two neighboring actin subunits, a region identified as an interface for many other actin-binding polypeptides, including Lifeact, the most widely used actin-binding probe. Thus, rather than contrasting F-tractin and Lifeact, our data indicate that these peptides have analogous modes of interaction with F-actin. Our study dissects the structural elements of F-tractin and provides a mechanistic basis for the selection and future development of actin probes.

## Introduction

Actin filaments (F-actin) are an essential component of the cell cytoskeleton, involved in numerous intracellular process, including cell movement, division and maintenance of cell shape^1^. Due to the critical role of F-actin in cellular morphogenesis, various actin probes have been developed to visualize actin under both physiological and pathological conditions^2^. These probes include small molecules^3, 4^, recombinant tags^5, 6^, actin-binding proteins^7, 8^ and peptides ^9,10^. However, the application of these compounds can disrupt normal cytoskeletal homeostasis, making it crucial to identify and understand the molecular basis of any side effects for accurate data interpretation.

Phalloidin, a cyclic peptide derived from the fungus *Amanita phalloides*, was one of the first probes used for F-actin labeling^11, 12^. This small molecule remains a gold standard for actin visualization due to its high affinity and selectivity for F-actin^2^. However, its low membrane permeability and stabilization of actin filaments limit its use to fixed cells^13^. The need to visualize actin dynamics led to the development of genetically encoded fluorescent probes. Chimeras of green fluorescent protein (GFP) with actin have been a simple and popular technique to visualize actin filaments in living cells^14, 15^. However, the bulky GFP tag on actin has been shown to significantly affect actin assembly^16, 17^. As alternatives to GFP-actin, GFP-labeled actin-binding domains such as from utrophin^7^ and fimbrin^18^ have been used. In addition, affimers and actin-binding nanobodies^8^ have been developed and successfully used in a variety of cell types and organisms.

To minimize steric clashes of large probes, small fluorophore-labeled peptides have been developed. The most commonly used is Lifeact, a 17-amino acid peptide derived from the N-terminus of the *Saccharomyces cerevisiae* actin-binding protein 140 (Abp140). This probe has been cited in over 1500 studies and in the majority of reports, Lifeact did not interfere with^9, 20, 21^ or had minimal effect on the cytoskeleton architecture^22, 23^. However, in certain experimental setups, Lifeact dramatically altered actin-dependent morphogenesis in yeast^24^, *Drosophila*^25^, zebrafish^26^, and mammalian cells^27^. A possible molecular mechanism for these cellular artifacts was proposed based on the cryo-EM structure of the Lifeact-F-actin complex^28, 29^. The structure and subsequent biochemical experiments demonstrated that Lifeact competes with cofilin, myosin and other actin binding factors for the same binding site on actin filaments, providing a possible basis for Lifeact-induced artifacts in cell morphogenesis. The availability of high-resolution structural data and the pressing need for better probes to visualize and control the actin cytoskeleton inspired the recent development of an optogenetic Lifeact variant^30^. In addition, the reports of Lifeact-induced artifacts motivated researchers to consider alternative actin-binding peptides as probes^31-34^.

F-tractin, a 43 amino acid-long peptide, is an alternative to Lifeact for the visualization of actin filaments^10, 35^. The peptide consists of residues 10–52 of the rat actin-binding inositol 1,4,5- trisphosphate 3-kinase A. F-tractin labels F-actin structures without causing female sterility or significant actin defects during *Drosophila* oogenesis^25^ and has been proven to reliably report on F-actin organization in primary neurons without notable effects on cell function^36^. However, in *Xenopus* XTC cells, expression of F-tractin induced radial actin bundles and longer filopodia^23^. Such bundling activity could also explain the reported increase in cell stiffness and reduction in cell migration associated with F-tractin^37^. Despite the widespread use of F-tractin, the molecular basis for these artifacts is poorly understood due to a lack of structural and biochemical data.

In this study, we determined the structure of the F-tractin-F-actin complex using cryo-EM. The 3.4 Å structure reveals that F-tractin is composed of a flexible N-terminal region and an amphipathic C-terminal helix. We show that the N-terminus is dispensable for F-actin binding, and its removal dramatically decreases actin bundling *in vitro* and increases the exchange rate in FRAP experiments. The C-terminal helix binds to a hydrophobic pocket formed by two actin subunits of the same filament strand. This binding site is similar to that of many actin-binding proteins, including cofilin and myosin, suggesting that F-tractin competes with them in cells. Our results help to predict potential experimental artifacts using F-tractin and provide a strong foundation for the development of improved actin-binding probes.

## Results and discussion

We produced the 43 amino acid F-tractin peptide using solid-phase synthesis. The peptide was nearly insoluble in water, Tris-buffered saline (TBS), or DMSO, but soluble in methanol to a concentration of 2 mM. Expecting a micromolar affinity of F-tractin for F-actin similar to Lifeact^28, 29^, we incubated preformed F-actin with 100 μM of the peptide to obtain a high decoration of actin filaments with the probe. The mixture was then applied onto a grid and plunge-frozen in liquid ethane. Cryo-EM analysis of the sample revealed large heterogeneous bundles instead of single actin filaments (Fig. 1a). These high-order structures impeded data analysis, prompting us to minimize the interaction time between F-actin and F-tractin. By mixing F-actin with the peptide immediately prior to application to the grid, we observed a sufficient number of individual actin filaments for data analysis. We collected and processed the dataset, resulting in a 3.4 Å reconstruction with local areas reaching 3 Å (Fig. 1b, S1, Supplementary Table 1). This reconstruction enabled us to build the atomic model of the F-tractin-F-actin complex.

**Fig. 1.**
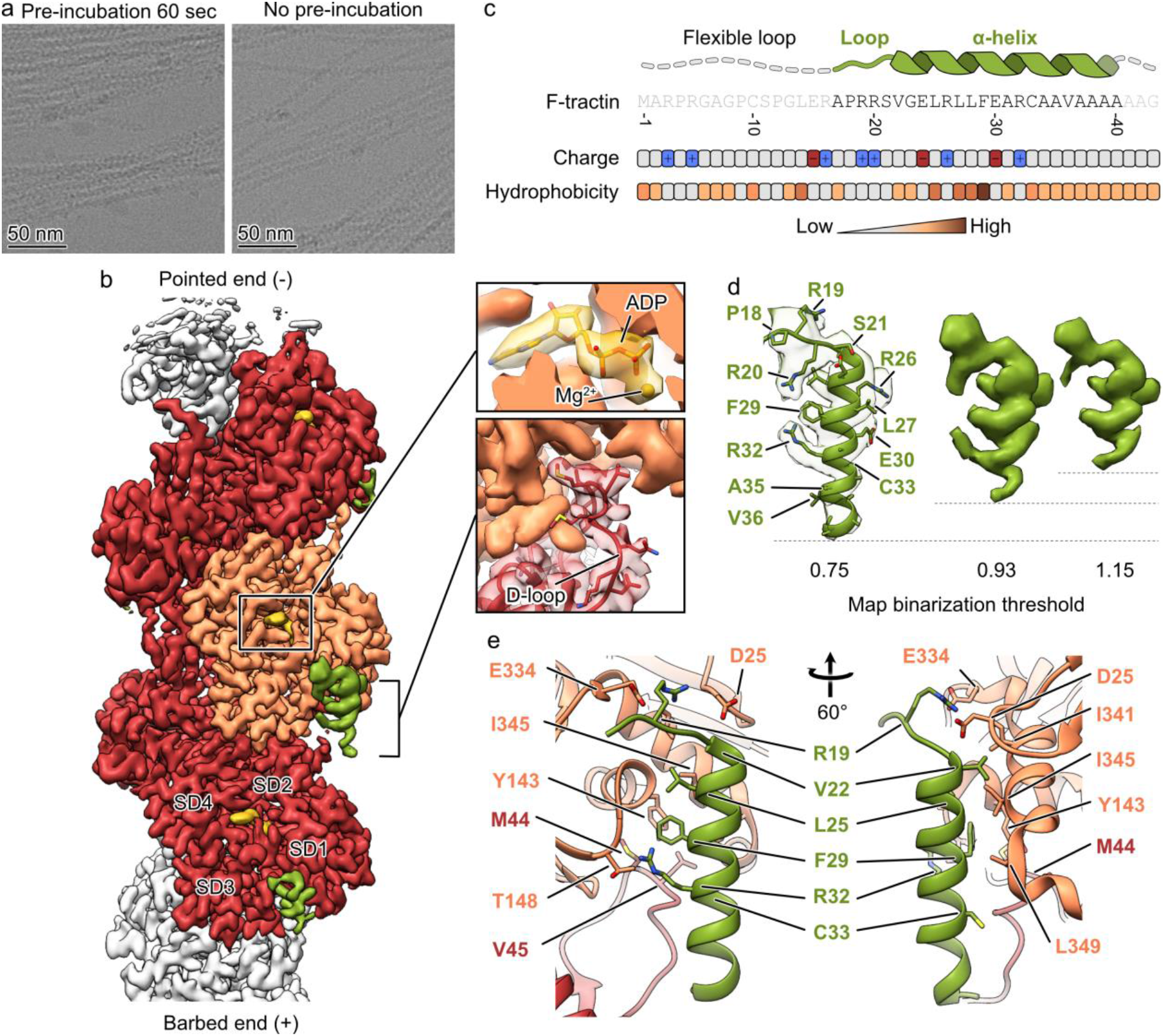
Cryo-EM structure of the F-tractin–F-actin complex. (a) Cryo-EM micrographs of actin filaments after mixing with F-tractin with (left) and without pre-incubation (right). (b) The 3.4 Å map of the F-tractin–F-actin complex shows a clear density for actin (orange and red), ADP (gold) and the F-tractin peptide (green). SD – subdomain. (c) The amino acid sequence of F-tractin, the distribution of its charged and hydrophobic residues and the schematic structure of the peptide. (d) F-tractin and its corresponding density at different map binarization thresholds, illustrating the flexibility of the extreme C-terminus. (e) Atomic model of the F- tractin-F-actin interface.

F-actin exhibited the expected ADP-state conformation with a closed D-loop (Fig. 1b). The RMSD values of 1.04 Å between the previously published structure of F-actin^41^ and F-actin in complex with F-tractin indicated that F-tractin did not alter the structure of F-actin. We were able to unambiguously build amino acids 17-40 of F-tractin (Fig. 1c and d). The 16 N-terminal residues, which include six prolines and glycines, could not be resolved due to their flexibility. Notably, this region also contains 3 positively charged arginines that can form non-specific interactions with negatively charged actin filaments, creating actin bundles.

Following the flexible region, the next five residues form a structured loop, with Arg-19 forming a salt bridge with Asp-25 and Glu-334 of actin. The peptide continues as an extended α-helix that spans two adjacent actin subunits along the same filament strand. The interaction interface between the helix and F-actin is primarily hydrophobic: Val-22, Leu-25, and Phe-29 of F-tractin fit into a hydrophobic pocket formed by Tyr-143, Ile-341, Ile-345, and Leu-349 of one actin subunit, and Met-44 and Val-45 of a neighboring subunit (Fig. 1e). Additionally, Arg- 32 forms hydrogen bonds with Thr-148 of actin, further stabilizing the complex. The adjacent Cys-33 faces the actin filament but does not engage in direct interactions. While Cys-33 may appear to be a potential site for covalent labeling with sulfhydryl-reactive groups such as maleimides, the close proximity of the attached chemical group to the actin filament could affect the properties of both the filament and the chemical group. Finally, the density quality of F-tractin decreases gradually toward the C-terminus (Fig. 1d), suggesting increased flexibility and the absence of specific interactions between the extreme C-terminal residues of F-tractin and F-actin.

The low solubility of F-tractin in aqueous solutions and its strong bundling effect on F-actin *in vitro* are significant disadvantages that prompted us to improve the peptide based on our structural data. We made three modifications: 1) we removed 16 N-terminal amino acids to eliminate potential nonspecific interactions with actin filaments; 2) we mutated cysteine 33 to alanine to prevent the formation of disulfide bonds that can lead to F-tractin oligomerization; 3) we substituted alanine 35 with serine, valine 36 with alanine, and removed the extreme C- terminus containing five alanines and one glycine to reduce the hydrophobicity of the C- terminal part of the peptide. The resulting optimized peptide, F-tractin_opt_, was chemically synthesized, water-soluble, and exhibited minimal actin bundling when used at concentrations up to 100 µM, as demonstrated by low-speed centrifugation assays and cryo-EM (Fig. 2b and c). These proof-of-principle experiments suggest that our structure of the F-tractin-F-actin complex provides a solid foundation for engineering improved actin visualization probes.

**Fig. 2.**
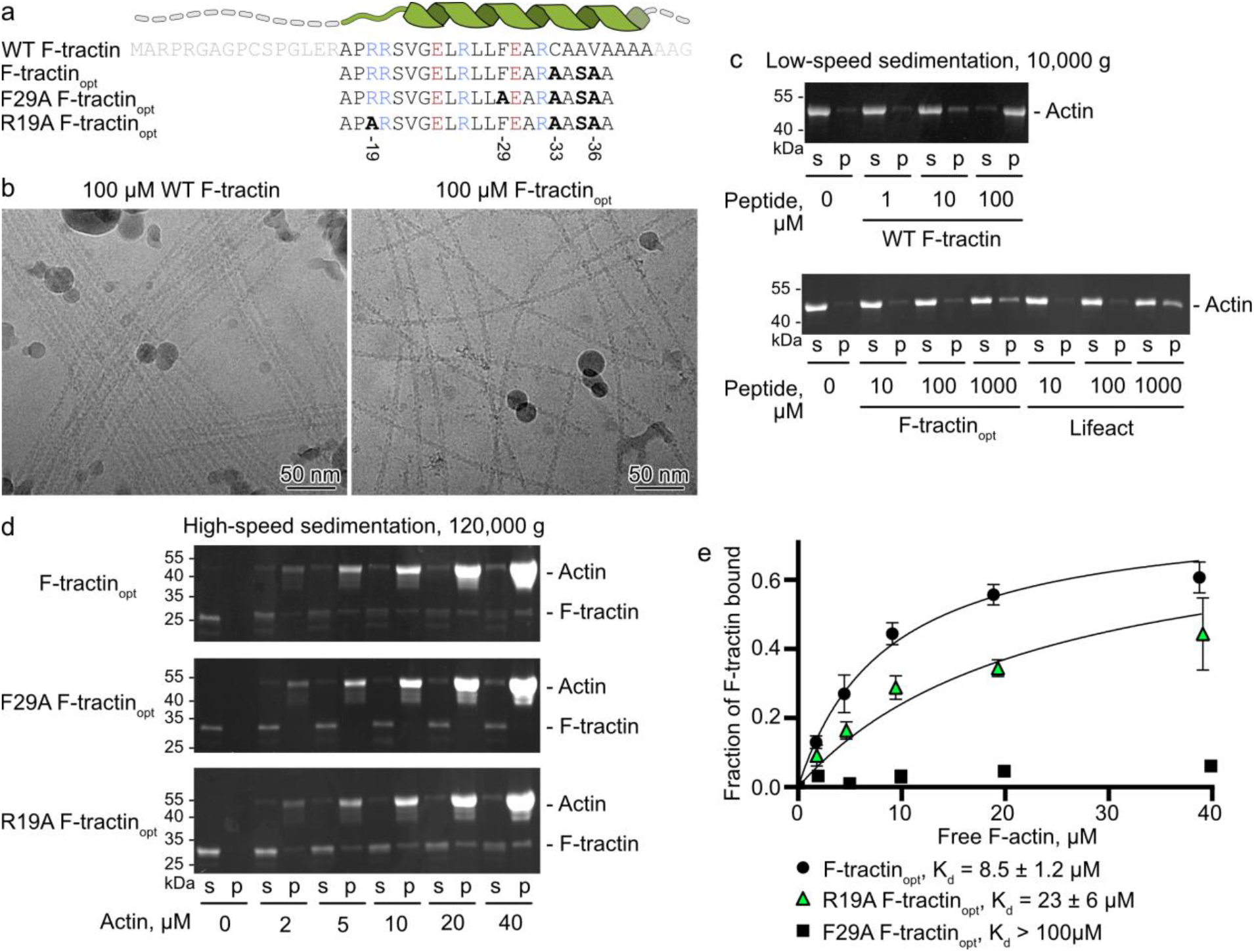
Point mutations and deletions in F-tractin alter its interaction to F-actin. (a) Amino acid sequence of WT F-tractin variants. Amino acids colored in light grey were not observed in the cryo-EM reconstruction. Amino acids colored in blue and red correspond to positively and negatively charged residues, respectively. (b) Cryo-EM micrographs of F-actin mixed with 100 µM of WT F-tractin or F-tractin_opt_. (c) Actin bundling assay using low-speed centrifugation of F-actin in the presence of WT F-Tractin, F-tractin_opt_, or Lifeact. Representative SDS-PAGE from 3 independent experiments are shown. (d) Cosedimentation of F-actin and 2 μM of F- tractin-mCherry with supernatant (s), and pellet (p) fractions analyzed by SDS-PAGE. The upper band corresponds to actin, and the lower band corresponds to F-tractin_opt_-mCherry. Representative stain-free gels are shown. (e) The fractions of F-tractin_opt_-mCherry that cosedimented with F-actin were quantified by densitometry and plotted against actin concentrations. Error bars in (e) correspond to standard deviations of 3 independent experiments. Uncropped gels can be found in Fig. S2 and data points for panel (e) can be found in Table S3.

In cell biology experiments, F-tractin is typically used in fusion with fluorescent proteins. Therefore, in order to estimate the interaction of F-tractin_opt_ with F-actin in a similar setup, we engineered and produced mCherry fusions with F-tractin_opt_ and tested their affinity for F-actin using high-speed cosedimentation assays (Fig. 2d and e). We observed a dose-dependent increase of F-tractin_opt_ in the pellet fraction and estimated the F-actin-F-tractin_opt_ K_d_ to be 8.5 ± 1.2 µM. Unfortunately, the production of mCherry fusion with WT F-tractin in *E. coli* resulted in an extremely low yield, preventing a direct comparison of the affinities between WT F- tractin-mCherry and F-tractin_opt_-mCherry. Nonetheless, the similar K_d_ of 6.8 µM observed for the longer 66-amino acid-long F-tractin fused to the carrier protein NusA^10^ suggests that our optimization did not greatly alter the F-tractin_opt_ affinity for F-actin. In agreement with the biochemical data, F-tractin-EGFP and F-tractin_opt_-EGFP showed no significant difference in actin labelling efficiency experiments in U-2 OS osteosarcoma cells (Fig. 3a, b and c). However, in fluorescence recovery after photobleaching (FRAP) experiments, F-tractin_opt_ showed faster exchange rates than WT F-tractin, suggesting higher exchange rates due to reduced interaction with actin (Fig. 3c, d and f).

**Fig. 3.**
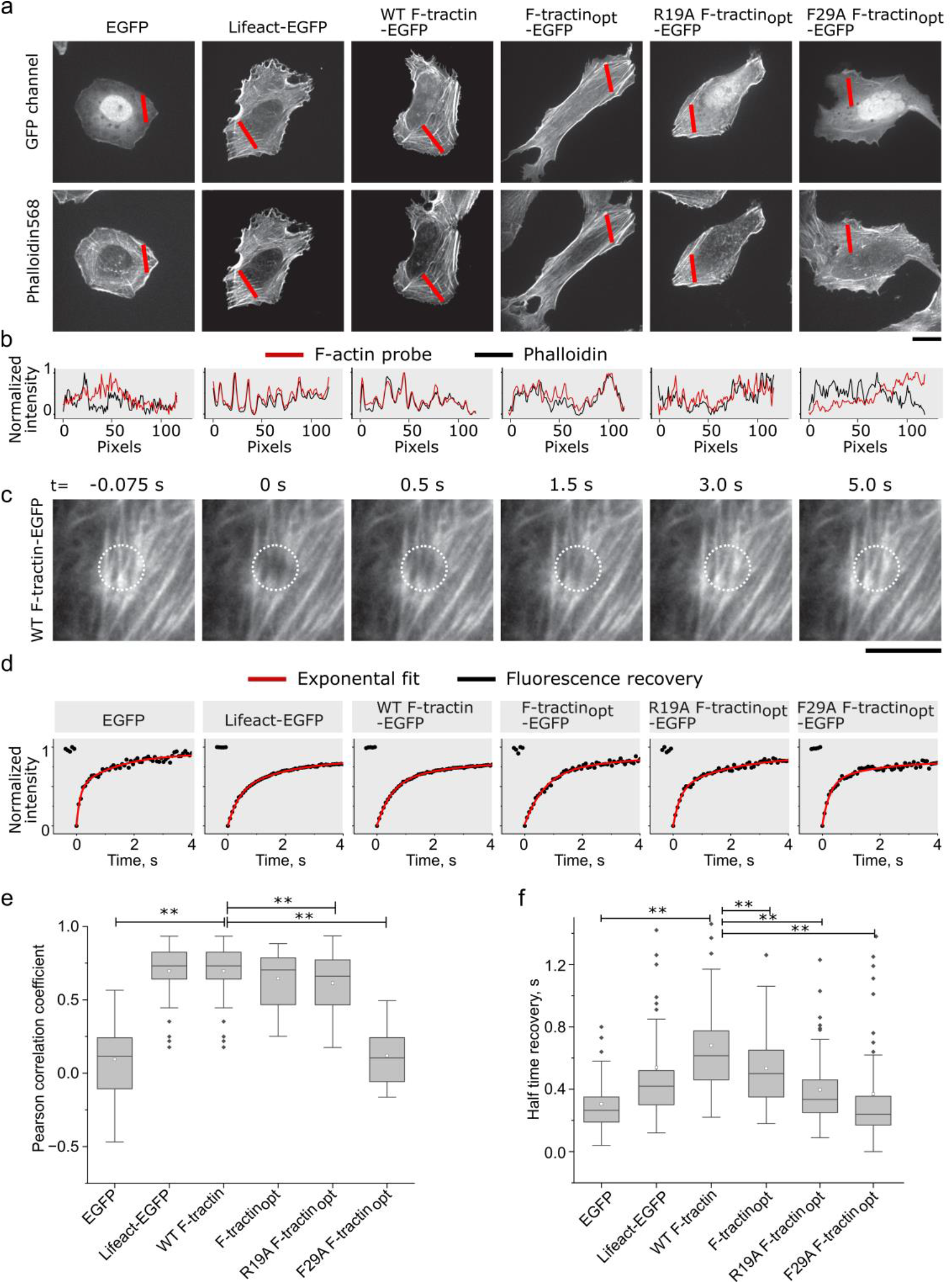
Actin labelling efficiency and exchange rate of F*-*tractin variants in U-2 OS cells. (a) Representative images of F-tractin-EGFP (GFP channel) and phalloidin. Scale bar: 20 μm. (b) Normalized line scans (lines indicated in red) of F-tractin variants (red) compared with respective phalloidin staining (black) (c) Time frames of a FRAP experiment in U2-OS cells expressing F-tractin-EGFP. Scale bar: 5 μm. (d) Normalized recovery profiles for indicated F- actin probes (black) and the respective exponential fits (red). (e) Boxplots of Pearson correlation coefficients between F-tractin variants and corresponding phalloidin staining. T-test p ≤ 0.01: ** (N = 3 experiments, n > 18 cells). (f) Boxplots of FRAP recovery half times for indicated F- tractin variants, free EGFP and Lifeact. T-test p ≤ 0.01: **. (N = 3 experiments, n > 25 cells). Data points for (e) and (f) in Table S3.

Having established this, we aimed to determine whether point mutations could further modify the F-tractin_opt_ interaction with F-actin. To this end, we generated two F-tractin_opt_ variants with the following mutations: Phe29Ala, to disrupt the central hydrophobic interaction with F-actin, and Arg19Ala, to eliminate the salt bridge between the central loop and negatively charged residues on the F-actin surface. Cosedimentation assays revealed that these mutants had significantly lower affinity for F-actin, with Phe29Ala F-tractin_opt_ being barely detectable in the pellet at the highest concentration of F-actin used (Fig. 2d and e). Consistent with the *in vitro* experiments, R19A F-tractin_opt_ exhibited a significant drop in colocalization with phalloidin in comparison to F-tractin_opt_. F29A F-tractin_opt_ completely failed to bind to actin structures in cells, (Fig. 3a, b and e). These results highlight the importance of the interactions formed by Arg-19 and Phe-29 of F-tractin_opt_ and demonstrate the potential of structure-based modifications to refine F-tractin interactions with F-actin.

Since its discovery, F-tractin has been considered an alternative to Lifeact for imaging cytoskeletal dynamics in living cells, but their comparison has been limited to cell biological studies^23, 25, 37^. Now, with the structures of both probes in complex with F-actin available, we can compare them in atomic details (Fig. 4a, b and c). Both probes are at their core amphipathic helices that bind to overlapping interfaces of consecutive actin subunits. Lifeact forms interactions with F-actin via five hydrophobic residues (Val-3, Leu-6, Ile-7, Phe-10, and Ile- 13), while F-tractin only utilizes three of these residues (Val-22, Leu-25, and Phe-29) and instead includes an additional N-terminal loop that forms salt bridges with Asp-25 and Glu-334 of actin. Interestingly, mutations of the phenylalanine (Phe-10 in Lifeact and Phe-29 in F-tractin) abolish the interactions of both probes with F-actin^28^ (Fig. 2e). They exhibit comparable *in vitro* affinities for F-actin (F-tractin_opt_: 8.5 ± 1.2 µM, Lifeact: 14.9 ± 1.6 μM^28^), cellular colocalization with phalloidin and FRAP recovery rates (Fig. 3). This suggests that both probes bind in an analogous fashion and consequently should also compete to a similar extent with other actin-binding proteins, such as cofilin, myosin, and bacterial effectors (Fig. 4d).

**Fig. 4.**
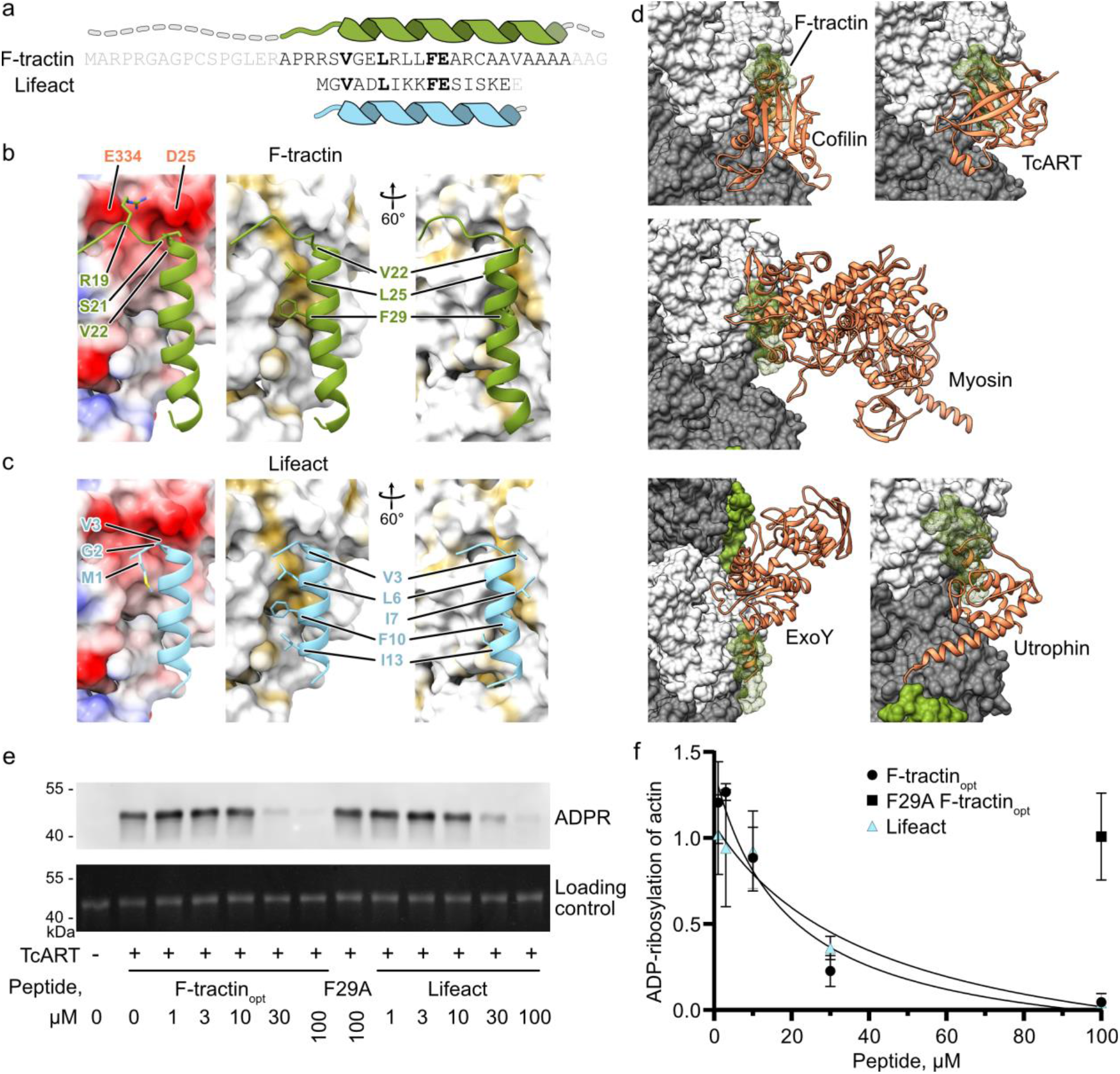
F-tractin and Lifeact share an interaction interface. (a) Comparison of amino acid sequence and secondary structure elements of F-tractin and Lifeact. (b, c) Comparison of electrostatic (left) and hydrophobic (middle/right) interactions of F-tractin and Lifeact (PDB 7AD9^28^). (d) Structures of cofilin (5YU8^40^), myosin (5JLH^41^), TcART (7Z7H^39^), ExoY (7P1G^42^) and utrophin (6M5G^29^) bound to F-actin (grey and white). Note that all proteins overlap the F-actin interaction site of F-tractin (green). (e) Levels of actin ADP-ribosylation by TcART in the presence of F-tractin_opt_ and Lifeact analyzed by western blot of a His-tagged ADP-ribose binding protein. (f) Level of ADP-ribosylation of actin were quantified by densitometry and plotted against peptide concentrations. Error bars in (f) correspond to standard deviations of 3 independent experiments. Uncropped gels can be found in Fig. S2 and data points for panel (f) can be found in Table S3.

To test this hypothesis, we produced the F-actin-binding bacterial ADP-ribosyltransferase effector TcART from *Photorhabdus luminescens*. This effector transiently interacts with F- actin at the F-tractin/Lifeact binding site^39^ and covalently modifies F-actin with easily detectable ADP-ribose, making it a model protein for competition assays with actin-binding peptides. We then used TcART to modify F-actin in the presence of different concentrations of Lifeact and F-tractin_opt_ (Fig. 4e and f). While Phe29Ala F-tractin_opt_, did not interfere with TcART modification of F-actin, both Lifeact and F-tractin_opt_ inhibited TcART activity in a similar dose-dependent manner. Thus, this experiment, together with structural and biochemical data, suggests that Lifeact and F-tractin_opt_ are nearly identical actin-visualizing probes and that the previously reported differences between Lifeact and F-tractin originate from the flexible distal areas of F-tractin or their expression levels, rather than from differences in their interactions with F-actin. These results provide cell biologists with crucial information for making informed decisions on the use of one or another actin-binding probe.

## Materials and methods

### Protein production and purification

Rabbit skeletal muscle α-actin was purified from muscle acetone powder (Pel-Freez Biologicals) as described in^38^ and stored in 50 µl aliquots at −70°C.

TcART, the ADP-ribosyltransferase from *P. luminescens* Tc toxin, was purified from *E. coli* BL21-CodonPlus(DE3)-RIPL harboring the plasmid pB656 as described previously^39^.

Three F-tractin variants fused to mCherry and the his-tagged ADP-ribose-binding protein Af1521 with mutations K35E and Y145R^43^ were purified from *E. coli* BL21-CodonPlus(DE3)- RIPL cells harboring the plasmids pB1043, 1044, 1045 and 791, respectively (Supplementary Table 2). A single colony was inoculated into 200 ml of LB media and grown at 37°C. At OD_600_ 0.6-1.0, protein expression was induced by the addition of IPTG to a final concentration of 0.02 mM. After 16 hours of expression at 22°C, the cells were harvested by centrifugation, resuspended in TBS (20 mM Tris pH 8 and 150 mM NaCl), and lysed by sonication. The soluble fraction was applied on TBS-equilibrated Protino Ni-IDA resin (Macherey-Nagel, Germany), washed, eluted with TBS supplemented with 250 mM imidazole, dialyzed against TBS and stored at −20°C.

WT F-tractin (MARPRGAGPCSPGLERAPRRSVGELRLLFEARCAAVAAAAAAG), F- tractin_opt_ (APRRSVGELRLLFEARAASAA) and F29A F-tractin_opt_ (APRRSVGELRLL**A**EARAASAA) peptides were synthesized by Genosphere, France, with > 95% purity.

### Cosedimentation assays

Freshly thawed mammalian β-actin was spun down at 120,000 g for 20 min at 4°C using a TLA-120.1 rotor, and the supernatant containing G-actin was collected. The protein was then polymerized by incubation in F buffer (120 mM KCl, 20 mM Tris pH 8, 2 mM MgCl_2_, 1 mM DTT and 1 mM ATP) for 2 hours on ice. Cosedimentation assays were performed in 20 μl volumes by first incubating F-actin with F-tractin_opt_-mCherry fusions for 5 minutes at room temperature, then centrifuging at 120,000 g using the TLA-120.1 rotor for 20 minutes at 4°C. After centrifugation, aliquots of the supernatant and resuspended pellet fractions were separated on SDS-polyacrylamide gels with 2,2,2-trichloroethanol^44^, visualized under ultraviolet light, and analyzed by densitometry using ImageJ. The Kd values were calculated using GraphPad Prism software.

### ADP-ribosylation by TcART

A total of 8-μl mixtures of 2 μg (4.8 μM final concentration) actin and actin-binding peptides at specified concentrations were preincubated for 5 minutes at room temperature in the buffer containing 1 mM NAD, 20 mM Tris pH 8, 150 mM NaCl, and 1 mM MgCl_2_. The ADP-ribosylation reaction was initiated by the addition of 0.2 ng (314 pM) of TcART to the mixture. After 10 minutes of incubation at 37°C, the reaction was stopped by adding Laemmli sample buffer and heating the sample at 95°C for 5 minutes. The components of the mixture were separated by SDS-PAGE, blotted onto a polyvinylidene difluoride (PVDF) membrane using a Trans-Blot Turbo Transfer System (Bio-Rad), and visualized using a combination of the His-tagged ADP-ribose binding protein at the concentration of 0.01 mg/ml and HisProbe-HRP conjugate. The level of actin ADP-ribosylation was quantified by densitometry using ImageJ.

### Actin-bundling assays

8 μl of actin filaments at 10 μM were mixed with peptides at the specified concentration, incubated for 5 min at room temperature, and spun down at 10,000 g for 10 min at 4°C. Aliquots of the supernatant and resuspended pellet fractions were separated on SDS-polyacrylamide gels with 2,2,2-trichloroethanol^44^ and visualized with ultraviolet light.

### Cryo-EM, data analysis, model building

Actin filaments were prepared as for cosedimentation assays. Shortly before plunging, F-actin was diluted to 6 µM with F-buffer. 18 µl of 6 µM F-actin was mixed with 1 µl of 2 mM WT F-tractin peptide in methanol or 2 mM F-tractin_opt_ in water and supplemented with 1 µl of 0.4% (w/v) of Tween-20 to improve ice quality. 3 µl of sample was applied onto a freshly glow-discharged copper R2/1 300 mesh grid (Quantifoil), blotted for 8 s on both sides with blotting force -15 and plunge-frozen in liquid ethane using the Vitrobot Mark IV system (Thermo Fisher Scientific) at 13 °C and 100% humidity.

The dataset was collected using a Talos Arctica transmission electron microscope (Thermo Fisher Scientific) equipped with an XFEG at 200 kV using the automated data-collection software EPU version 2.7 (Thermo Fisher Scientific). 2 images per hole with defocus range of -0.5 - -2.5 µm were collected with a K3 detector (Gatan) operated in super-resolution mode. Image stacks with 50 frames were collected with a total exposure time of 3.5 sec and a total dose of 45 e^-^/Å^2^. 8100 movies were collected and 4048 of them were used for data processing. After motion correction performed in Relion version 3.1, the micrographs were imported into Cryosparc for patch CTF correction. 657 micrographs were discarded due to excessive drift and suboptimal ice thickness. Filaments from the remaining 3391 micrographs were picked using Filament Tracer in Cryosparc with filament diameter 75 Å and separation distance between segments 52.5 Å. 1474077 particles with 256 px box size were extracted and 2D classified into 200 classes. 957326 particles were selected for Helix Refine with a helical twist estimate of 166.4° and a helical rise estimate of 27.6 Å. After this 3D refinement, a round of global and local CTF refinements and another helical refinement, the reconstruction reached 3.3 Å. To improve the density corresponding to the F-tractin peptide, alignment-free 3D classification with a focused mask was used to separate particles into 2 classes. The larger class containing the density for the F-tractin peptide with 502568 particles was locally refined with the ‘use pose/shift gaussian prior during alignment’ setting to allow only small rotations and shifts of the reconstruction. The B-factor sharpened output of the local refinement was used for model building.

The structure of F-tractin, predicted in Alphafold 2^45^, and the structure of ADP-F-actin^38^ (PDB 5ONV) were fitted into the density and refined in Isolde^46^ and Phenix^47^.

### Cell culture and Transfection

U2OS human osteosarcoma cells were cultured in McCoy’s medium (with 10% fetal calf serum (FCS)). Cells were seeded at a density of 30000 cells/cm^2^ in an 8-well plate (Sarstedt) and allowed to grow overnight at 37°C (10% CO_2_) prior to transfection^48^. The plasmids of all desired actin markers fused with GFP were transfected using the X-fect transfection reagent (Takara Bio) as per the manufacturer’s recommendation. 500 ng of each plasmid was mixed with X- fect reaction buffer to make up a final volume of 50 μl. 0.5 μl of Xfect Polymer was added and vortexed shortly before incubating the mixture at room temperature for 10 minutes. The mixture was spin-down and pipetted into the 8 well of the plate. The cells were then incubated at 37°C overnight before imaging.

### Fixing and staining

Cells were fixed and stained with Phalloidin-568 (Promokine). The cell culture medium (McCoy’s) was discarded, and cells were washed once with Phosphate-Buffered Saline (PBS). The cells were fixed with 4% paraformaldehyde (PFA) (50 ul per well) for 20-30 minutes at room temperature and washed twice with PBS. 0.01% Triton X-100 (TX-100) was used to permeate cells (3 to 5 minutes at RT), followed by a two-time PBS washing. The cells were incubated at room temperature with 100 nM phalloidin for 20 minutes and then washed twice with PBS. Cells were then stained with Hoechst (1:5000 dilution from the original stock (Thermo Fischer, H3570)) for 15 minutes, followed by two times of PBS washing.

### Imaging

We imaged the fixed cells with the iMIC system from FEI/Till Photonics equipped with a spinning disk unit (Andromeda), using a 40X lens (0.9 NA, Olympus) and lasers at wavelengths of 405 nm, 488 nm and 561 nm (Omicron Sole-6). Multichannel images were captured using an EMCCD camera (Andor iXon Ultra 897) controlled by Live Acquisition software (Till Photonics). For photobleaching experiments, we used another iMIC system from FEI/Till Photonics, for capturing videos with an Olympus 100× oil immersion lens (1.4 NA). The TIRF angles were adjusted prior to each microscopy session. We used 488 nm (Cobolt Calypso, 75 mW) to excite the EGFP fused F-tractin constructs in cells. For each cell that was imaged, there were 3-4 FRAP Region of Interests (ROI) (100% laser for 100 ms) and a Normalization ROI (0% laser 100 ms). Movies were acquired with frame interval 76 ms and ROIs were bleached sequentially after 100 frames. The timelapse videos were captured Imago-QE Sensicam camera.

### Quantification of recovery rates from FRAP movies

The FRAP videos were imported into Fiji for analysis. Point FRAP regions of interest (ROIs) were expanded to circles with a radius of 20 pixels. We customized the standard Fiji FRAP plugin to obtain the altered FRAP ROIs and the normalizing ROI. The plugin measures the average intensity values for each ROI over a specified number of image slices. We then normalize these intensity values, determine the bleaching frame, and calculate the mean intensity before bleaching. The plugin then fits the normalized FRAP recovery curve using an exponential recovery model and calculates half time recovery. We used a single term exponential function as reported in previous studies (Koskinen Frontiers in neuroanatomy 2014). The half time recovery values of different variants were compared pairwise to WT F-tractin using Student t-test.

### Quantification of Pearson Correlation coefficient

From the multichannel images captured using the spinning disk microscope, we segmented the cells and nuclei automatically using Cellpose. We used the standard Cyto2 and nuclei modules of Cellpose for generating cell and nuclear masks. Following that, we subtracted the nucleus from the cell mask and performed the correlation between the EGFP and Phalloidin slices of the image. We used *pearsonr* function from the scipy.stats module to calculate the Pearson correlation coefficient and the associated p-value for testing correlation. We used Student t-test for the statistical comparison of dataset.

## Acknowledgements

We thank Artem Stetsenko, Roman Koning and Frank Faas for help with cryo-EM data collection and Michiel Punter for maintaining the computing cluster. Cryo-EM data were collected at the electron microscopy facilities of the University of Groningen and Leiden Medical Center. This work has been funded by the University of Groningen (AB) and the German Research foundation (DFG, SFB1009, SFB1348 and SFB1557 to RWS).

## Author contribution

A.B. and R.W.-S. designed and supervised the project. A.B. and D.S. prepared cryo-EM specimens, collected and analyzed EM data. D.S. and A.B. performed and analyzed biochemical experiments. A.S. performed cell biology experiments and analyzed the data. All authors contributed to preparing figures and writing the manuscript.

## Data availability

The coordinates for the cryo-EM structure of the F-tractin-F-actin complex have been deposited in the Electron Microscopy Data Bank under accession numbers EMD-51496. The corresponding molecular model have been deposited at the wwPDB with accession code PDB 9GOB. The raw data generated during the current study are available from the corresponding authors on reasonable request. Individual data points from the graphs at Fig. 2, 3 and 4 are available in Supplementary Table 3. Uncropped gels and western blots can be found in Supplementary Fig. 2.

