## Supplementary Table 1 for "Structure of the F-tractin–F-actin complex"

Supplementary information

**Supplementary Table 1. Cryo-EM data collection, refinement, and validation statistics**

| Microscope | Talos Arctica |
| --- | --- |
| Voltage (kV) | 200 |
| Defocus range (µm) | -0.5 to -2.5 |
| Camera | Gatan K3 Superresolution mode |
| Pixel size (Å) | 1.04 |
| Total electron dose (e/Å^2^) | 45 |
| Exposure time (s) | 3.5 |
| Frames per movie | 50 |
| Number of movies | 3391 (8100) |
| **3D Refinement** | |
| Number of particles | 502568 |
| Final resolution (Å) | 3.4 |
| Helical rise (Å) | 28.9 |
| Helical twist (°) | -166.7 |
| **Atomic model statistics** | |
| Non-hydrogen atoms | 15660 |
| Molprobity score | 1.54 |
| Clashscore | 4.92 |
| Bond RMSD (Å) | 0.003 |
| Angle RMSD (°) | 0.735 |
| Poor rotamers (%) | 0 |
| Favored rotamers (%) | 99.63 |
| Ramachandran favored (%) | 95.93 |
| Ramachandran allowed (%) | 4.07 |
| Ramachandran outliers (%) | 0 |

**Supplementary Table 2. List of primers and plasmids used in this study.**

| **Plasmid** | **Description** | **Reference** |
| --- | --- | --- |
| pB791 pET28a eAF1521 | Gene, encoding ADP-ribose binding protein AF 1521 (uniprot O28751) with two mutations K35E and Y145R (Kathrin Nowak Nat Commun 2020), was synthesized by Twist Biosciences and cloned into pET28a vector using NdeI/NotI sites. | This study |
| pB1030 pET28a F-tractin | Gene, encoding F-tractin in fusion with mCherry-6xHis, was synthesized by Twist Biosciences and cloned into pET28a vector using NcoI/HindIII. | This study |
| pB1043 pET28a F-tractin_opt_ | Primers catggccccacgacgttcagtcggggagctcagattgctttttgaagcgcgggcagcatctgccgccg and gatccggcggcagatgctgcccgcgcttcaaaaagcaatctgagctccccgactgaacgtcgtggggc were annealed to generate DNA sequence encoding F-tractin_opt_ and inserted into pB1030 digested with NcoI and BamHI. | This study |
| pB1044 pET28a F-tractin_opt_ F29A | Primers catggccccacgacgttcagtcggggagctcagattgcttgcggaagcgcgggcagcatctgccgccg and gatccggcggcagatgctgcccgcgcttccgcaagcaatctgagctccccgactgaacgtcgtggggc were annealed to generate DNA sequence encoding F29A F-tractin_opt_ and inserted into pB1030 digested with NcoI and BamHI. | This study |
| pB1045 pET28a F-tractin_opt_ R19A | Primers catggccccatcacgttcagtcggggagctcagattgctttttgaagcgcgggcagcatctgccgccg and gatccggcggcagatgctgcccgcgcttcaaaaagcaatctgagctccccgactgaacgtgatggggc were annealed to generate DNA sequence encoding R19A F-tractin_opt_ and inserted into pB1030 digested with NcoI and BamHI. | This study |
| pB656 pET28 MBP TcART | His-tagged TcART in MBP fusion. | [1] |
| pB1055 pEGFP-N1 | Vector for expression of EGFP fusion genes in eukaryotic hosts. | Addgene plasmid # 13031 |
| pB1057 pEGFP-N1 F-tractin | Primers tatactcgagatggctcgcccgagaggggctggtccatgcagcccgggtttagagcgggccccacgacgttcagtcggggagctc and tataggatcccctgcggccgctgctgcggctacggctgcgcaccgcgcttcaaaaagcaatctgagctccccgactgaacgtcg were annealed and PCR-amplified to generate DNA sequence encoding F-tractin_,_ cut with XhoI and BamHI and inserted into pB1055 digested with XhoI and BamHI. | This study |
| pB1058 pEGFP-N1 F-tractin_opt_ | Primers tcgagatgGCCCCACGACGTTCAGTCGGGGAgCTcAGATTGCTTTTTGAAGCGCGGGCAGCAtctGCCGCCggg and gatccccGGCGGCagaTGCTGCCCGCGCTTCAAAAAGCAATCTgAGcTCCCCGACTGAACGTCGTGGGGCcatc were annealed to generate DNA sequence encoding F-tractin_opt_ and inserted into pB1055 digested with XhoI and BamHI. | This study |
| pB1059 pEGFP-N1 F-tractin_opt_ F29A | Primers tcgagatgGCCCCACGACGTTCAGTCGGGGAgCTcAGATTGCTTGCGGAAGCGCGGGCAGCAtctGCCGCCggg and gatccccGGCGGCagaTGCTGCCCGCGCTTCCGCAAGCAATCTgAGcTCCCCGACTGAACGTCGTGGGGCcatc were annealed to generate DNA sequence encoding F29A F-tractin_opt_ and inserted into pB1055 digested with XhoI and BamHI. | This study |
| pB1060_pEGFP-N1 F-tractin_opt_ R19A | Primers tcgagatgGCCCCATCACGTTCAGTCGGGGAgCTcAGATTGCTTTTTGAAGCGCGGGCAGCAtctGCCGCCggg and gatccccGGCGGCagaTGCTGCCCGCGCTTCAAAAAGCAATCTgAGcTCCCCGACTGAACGTGATGGGGCcatc were annealed to generate DNA sequence encoding R19A F-tractin_opt_ and inserted into pB1055 digested with XhoI and BamHI. | This study |
| pEGFP-N1 Lifeact | Primers aagcttcgaattcATGGGTGTCGCAGATTTGATCAAGAAATTCGAAAGCATCTCAAAGGAAGAAggg and tcgaattcATGGGTGTCGCAGATTTGATCAAGAAATTCGAAAGCATCTCAAAGGAAGAAggggatc were annealed to generate DNA sequence encoding Lifeact and inserted to pB1055 with BamHI and HindIII | This  study |

**Supplementary Table 3. Data points**

**Figure 2e**

|  | Fraction of **F-tractin_opt_** bound | | |
| --- | --- | --- | --- |
| F-actin, µM | Repeat 1 | Repeat 2 | Repeat 3 |
| 2 | 0,113982 | 0,125633 | 0,150215 |
| 5 | 0,211009 | 0,27957 | 0,318841 |
| 10 | 0,480176 | 0,433657 | 0,418605 |
| 20 | 0,569748 | 0,578018 | 0,523474 |
| 40 | 0,557282 | 0,621935 | 0,643478 |

|  | Fraction of F29A **F-tractin_opt_** bound | | |
| --- | --- | --- | --- |
| F-actin, µM | Repeat 1 | Repeat 2 | Repeat 3 |
| 2 | 0,02963 | 0,031317 | 0,036 |
| 5 | 0,020561 | 0,008869 | 0,001393 |
| 10 | 0,051546 | 0,016783 | 0,023887 |
| 20 | 0,035971 | 0,051087 | 0,050633 |
| 40 | 0,071269 | 0,046373 | 0,065217 |

|  | Fraction of R19A **F-tractin_opt_** bound | | |
| --- | --- | --- | --- |
| F-actin, µM | Repeat 1 | Repeat 2 | Repeat 3 |
| 2 | 0,078051 | 0,069519 | 0,125538 |
| 5 | 0,150979 | 0,148265 | 0,193573 |
| 10 | 0,31152 | 0,304965 | 0,249039 |
| 20 | 0,358879 | 0,357945 | 0,31634 |
| 40 | 0,488372 | 0,519149 | 0,323824 |

**Figure 3e**

| Pearson Correlation coefficient for each cell analyzed | | | | | |
| --- | --- | --- | --- | --- | --- |
| EGFP | Lifeact | WT F- tractin | F-tractin_opt_ | F29A F-tractin_opt_ | R19A F-tractin_opt_ |
| 0.10954 | 0.76643 | 0.78651 | 0.67295 | 0.30638 | 0.52439 |
| 0.17413 | 0.35274 | 0.83482 | 0.4789 | 0.35424 | 0.51659 |
| 0.20843 | 0.89957 | 0.81213 | 0.43512 | 0.05418 | 0.21147 |
| 0.56273 | 0.79351 | 0.73699 | 0.71914 | -0.09177 | 0.45312 |
| 0.25628 |  | 0.81463 | 0.24809 | 0.24243 | 0.58155 |
| -0.11892 | 0.76541 | 0.51708 | 0.17482 | 0.33315 | 0.34979 |
| -0.14581 | 0.75694 | 0.70416 | 0.74647 | -0.1638 | 0.21057 |
| -0.24725 | 0.63211 | 0.425 | 0.81435 | 0.10337 | 0.49853 |
| -0.06922 | 0.6686 | 0.41105 | 0.72387 | 0.49504 | 0.52201 |
| -0.33835 | 0.44503 | 0.6836 | 0.90307 | 0.07811 | -0.00839 |
| -0.35467 | 0.64123 | 0.65265 | 0.51485 | 0.00323 | 0.47492 |
| 0.18496 | 0.2489 | 0.46728 | 0.77871 | -0.05819 | 0.57243 |
| -0.46832 | 0.76761 | 0.86762 | 0.78517 | 0.24261 | 0.41911 |
| -0.02363 | 0.72541 | 0.84982 | 0.30793 | -0.06018 | 0.46935 |
| 0.12068 | 0.6592 | 0.30961 | 0.45293 | 0.1486 | 0.53496 |
| 0.56515 | 0.69321 | 0.67704 | 0.571 | 0.152 | 0.08256 |
| 0.0863 | 0.9123 | 0.68198 | 0.65055 | -0.10862 | 0.48473 |
| 0.45723 | 0.834 | 0.34589 | 0.76628 |  | -0.15254 |
| 0.44998 | 0.87403 | 0.46251 | 0.37407 |  | 0.52933 |
| 0.31887 | 0.70696 | 0.71653 | 0.53954 |  | 0.26181 |
| 0.0651 | 0.73191 | 0.81744 | 0.70429 |  | 0.72892 |
| 0.54355 | 0.4622 | 0.82248 | 0.62755 |  | 0.44463 |
| -0.13263 | 0.21844 | 0.78101 | 0.80792 |  | 0.56268 |
| 0.17658 | 0.78344 | 0.60454 | 0.9354 |  | 0.44615 |
| 0.06247 | 0.89871 | 0.25161 |  |  | 0.24766 |
| 0.2267 | 0.76498 | 0.64922 |  |  | 0.16194 |
| -0.22143 | 0.69642 | 0.44011 |  |  | -0.06174 |
| 0.34798 | 0.85502 | 0.70425 |  |  | 0.01029 |
| 0.22207 | 0.70383 | 0.81156 |  |  | 0.67663 |
| -0.09331 | 0.78924 | 0.63599 |  |  | 0.79555 |
| 0.15295 | 0.17659 | 0.88315 |  |  | 0.4683 |
| 0.17203 | 0.71164 | 0.7852 |  |  |  |
| 0.0401 | 0.93357 | 0.40483 |  |  |  |
| 0.01924 | 0.61379 | 0.73442 |  |  |  |
| -0.43342 | 0.82511 | 0.72687 |  |  |  |
| 0.49162 | 0.64172 | 0.78591 |  |  |  |
| 0.10954 | 0.91147 | 0.28591 |  |  |  |
| 0.17413 | 0.88287 |  |  |  |  |
| 0.20843 |  |  |  |  |  |
| 0.56273 |  |  |  |  |  |
| 0.25628 |  |  |  |  |  |
| -0.11892 |  |  |  |  |  |
| -0.14581 |  |  |  |  |  |
| -0.24725 |  |  |  |  |  |
| -0.06922 |  |  |  |  |  |
| -0.33835 |  |  |  |  |  |
| -0.35467 |  |  |  |  |  |
| 0.18496 |  |  |  |  |  |
| -0.46832 |  |  |  |  |  |
| -0.02363 |  |  |  |  |  |
| 0.12068 |  |  |  |  |  |
| 0.56515 |  |  |  |  |  |
| 0.0863 |  |  |  |  |  |
| 0.45723 |  |  |  |  |  |
| 0.44998 |  |  |  |  |  |
| 0.31887 |  |  |  |  |  |
| 0.0651 |  |  |  |  |  |
| 0.54355 |  |  |  |  |  |
| -0.13263 |  |  |  |  |  |
| 0.17658 |  |  |  |  |  |
| 0.06247 |  |  |  |  |  |
| 0.2267 |  |  |  |  |  |
| -0.22143 |  |  |  |  |  |
| 0.34798 |  |  |  |  |  |
| 0.22207 |  |  |  |  |  |
| -0.09331 |  |  |  |  |  |
| 0.15295 |  |  |  |  |  |
| 0.17203 |  |  |  |  |  |
| 0.0401 |  |  |  |  |  |
| 0.01924 |  |  |  |  |  |
| -0.43342 |  |  |  |  |  |
| 0.49162 |  |  |  |  |  |

**Figure 3f**

| Half time recovery – For each photobleached ROI | | | | | |
| --- | --- | --- | --- | --- | --- |
| EGFP | Lifeact | WT-tractin | F-tractin_opt_ | F29A F-tractin_opt_ | R19A F-tractin_opt_ |
| 0.09 | 0.28 | 0.55 | 0.74 | 0.13 | 0.5 |
| 0.19 | 0.3 | 0.51 | 0.7 | 0.12 | 0.32 |
| 0.22 | 0.37 | 0.47 | 0.61 | 0.22 | 0.23 |
| 0.29 | 0.29 | 0.3 | 0.68 | 0.23 | 0.31 |
| 0.06 | 0.19 | 0.43 | 0.24 | 0.19 | 0.27 |
| 0.16 | 0.18 | 0.76 | 0.47 | 0.62 | 0.17 |
| 0.11 | 0.52 | 0.8 | 0.67 | 0.12 | 0.27 |
| 0.28 | 1.2 | 0.71 | 0.56 | 1.25 | 0.33 |
| 0.04 | 0.44 | 0.9 | 0.41 | 0.24 | 0.25 |
| 0.15 | 0.34 | 0.53 | 0.73 | 0.2 | 0.21 |
| 0.17 | 0.26 | 0.77 | 0.38 | 1.19 | 0.21 |
| 0.22 | 0.52 | 0.46 | 0.34 | 0.48 | 0.28 |
| 0.25 | 0.27 | 0.86 | 0.47 | 0.17 | 0.29 |
| 0.23 | 0.26 | 0.68 | 0.31 | 0.17 | 0.54 |
| 0.44 | 0.44 | 0.9 | 0.29 | 0.43 | 0.28 |
| 0.15 | 0.52 | 0.77 | 1.26 | 0.2 | 0.31 |
| 0.07 | 0.62 | 0.98 | 0.35 | 0.16 | 0.29 |
| 0.12 | 0.41 | 0.61 | 0.64 | 0.31 | 0.39 |
| 0.19 | 0.4 | 0.42 | 0.35 | 0.17 | 0.19 |
| 0.14 | 0.8 | 0.46 | 0.35 | 0.16 | 0.25 |
| 0.05 | 0.26 | 0.91 | 0.61 | 0.14 | 0.32 |
| 0.16 | 0.16 | 0.74 | 0.54 | 0.11 | 0.2 |
| 0.14 | 0.44 | 0.96 | 0.57 | 0.28 | 0.53 |
| 0.14 | 0.53 | 0.39 | 0.52 | 0.54 | 0.33 |
| 0.12 | 0.57 | 0.48 | 0.22 | 0.11 | 0.17 |
| 0.16 | 0.42 | 0.35 | 0.3 | 0.06 | 0.39 |
| 0.26 | 0.48 | 0.76 | 0.41 | 0.15 | 0.43 |
| 0.2 | 0.37 | 0.49 | 0.29 | 1.11 | 0.86 |
| 0.4 | 0.22 | 0.37 | 0.27 | 0.09 | 0.24 |
| 0.17 | 0.25 | 1.04 | 0.65 | 0.19 | 0.25 |
| 0.13 | 0.33 | 2.85 | 0.43 | 0.31 | 0.59 |
| 0.14 | 0.35 | 0.59 | 0.74 | 0.31 | 0.25 |
| 0.25 | 0.48 | 1.27 | 0.3 | 0.5 | 0.18 |
| 0.13 | 0.27 | 0.75 | 0.57 | 0.13 | 0.31 |
| 0.31 | 0.26 | 0.88 | 0.4 | 0.21 | 0.3 |
| 0.57 | 0.35 | 0.51 | 0.41 | 0.39 | 0.29 |
| 0.08 | 0.27 | 0.74 | 0.65 | 0.13 | 0.55 |
| 0.04 | 0.31 | 0.69 | 0.43 | 0.25 | 0.36 |
| 0.27 | 0.52 | 0.66 | 0.49 | 0.26 | 0.22 |
| 0.4 | 0.77 | 0.67 | 0.57 | 0.3 | 0.24 |
| 0.22 | 0.48 | 0.67 | 0.69 | 0.18 | 1.23 |
| 0.18 | 0.16 | 0.32 | 0.54 | 0.3 | 0.38 |
| 0.3 | 0.57 | 0.4 | 1 | 0.31 | 0.31 |
| 0.41 | 0.32 | 0.71 | 0.47 | 0.47 | 0.34 |
| 0.19 | 0.5 | 1.17 | 0.28 | 0.19 | 0.24 |
| 0.43 | 0.85 | 1.7 | 0.47 | 0.76 | 0.25 |
| 0.52 | 0.32 | 0.58 | 0.42 | 0.41 | 0.23 |
| 0.27 | 0.44 | 0.48 | 0.43 | 0.46 | 0.37 |
| 0.27 | 0.44 | 0.22 | 1.26 | 0.31 | 0.55 |
| 0.3 | 0.48 | 0.6 | 0.34 | 0.16 | 0.62 |
| 0.48 | 0.31 | 0.73 | 1.01 | 2.72 | 0.64 |
| 0.38 | 0.42 | 0.48 | 0.86 | 0.46 | 0.49 |
| 0.34 | 0.36 | 0.56 | 0.33 | 0.23 | 0.19 |
| 0.17 | 0.29 | 0.78 | 0.19 | 0.24 | 0.31 |
| 0.16 | 0.39 | 0.42 | 0.48 | 0.42 | 0.27 |
| 0.43 | 0.45 | 0.53 | 0.45 | 0.7 | 0.46 |
| 0.41 | 0.69 | 1.46 | 0.6 | 0.16 | 0.26 |
| 0.31 | 0.36 | 0.41 | 0.35 | 0.26 | 0.34 |
| 0.25 | 0.57 | 0.45 | 0.56 | 0.31 | 0.36 |
| 0.26 | 0.69 | 0.75 | 0.24 | 0.44 | 0.43 |
| 0.27 | 0.12 | 0.57 | 0.71 | 0.22 | 0.37 |
| 0.8 | 0.45 | 0.84 | 0.68 | 0.12 | 0.31 |
| 2.01 | 0.19 | 0.91 | 0.8 | 0.28 | 0.41 |
| 0.4 | 0.61 | 0.59 | 0.48 | 0.42 | 0.41 |
| 0.14 | 0.3 | 0.46 | 1 | 0.23 | 0.21 |
| 0.23 | 0.43 | 1.04 | 1.06 | 0.13 | 0.55 |
| 0.34 | 0.91 | 0.38 | 0.42 | 0.45 | 0.69 |
| 0.35 | 0.4 | 0.53 | 0.55 | 1.01 | 0.38 |
| 0.32 | 0.35 | 0.8 | 0.56 | 0.21 | 0.2 |
| 0.26 | 0.4 | 0.51 | 0.18 | 0.27 | 0.3 |
| 0.23 | 0.85 | 0.69 | 0.49 | 0.22 | 0.24 |
| 0.73 | 0.95 | 0.63 | 0.4 | 0.33 | 0.6 |
| 0.26 | 0.55 | 0.83 | 0.62 | 0.12 | 0.26 |
| 0.28 | 0.27 | 0.94 | 0.31 | 0.27 | 0.33 |
| 0.21 | 0.15 | 0.75 | 0.27 | 0.3 | 0.23 |
| 0.5 | 0.24 | 1.93 | 0.37 | 0.39 | 0.24 |
| 1.84 | 0.4 | 0.29 | 0.19 | 0.23 | 0.46 |
| 0.17 | 0.4 | 0.44 | 0.19 | 0.15 | 0.37 |
| 0.58 | 0.48 | 0.66 | 0.2 | 0.2 | 0.11 |
| 0.31 | 0.44 | 0.39 | 0.38 | 0.26 | 0.47 |
| 0.23 | 0.43 | 1.37 | 0.18 | 0.3 | 0.3 |
| 0.52 | 0.68 | 0.4 | 0.65 | 0.34 | 0.46 |
| 0.41 | 0.5 | 0.43 | 0.35 | 0.14 | 0.17 |
| 0.64 | 0.3 | 0.55 | 0.57 | 0 | 0.78 |
| 0.23 | 1.56 | 0.71 | 0.43 | 0.24 | 0.19 |
| 0.32 | 0.69 | 0.81 | 0.53 | 0.23 | 0.24 |
| 0.24 | 7.88 | 0.72 | 0.44 | 0.43 | 0.25 |
| 0.42 | 0.49 | 0.29 | 0.25 | 0.11 | 0.4 |
| 0.35 | 0.14 | 0.99 | 0.64 | 0.25 | 0.37 |
| 0.28 | 0.58 | 0.46 | 0.54 | 0.64 | 0.4 |
| 0.28 | 0.47 | 0.77 | 0.53 | 0.28 | 0.22 |
| 0.19 | 0.38 | 0.3 | 0.52 | 0.33 | 0.72 |
| 0.36 | 0.41 | 0.68 | 0.29 | 0.24 | 0.35 |
| 0.45 | 0.39 | 1.08 | 0.96 | 0.18 | 0.42 |
| 0.25 | 0.41 | 0.62 | 0.78 | 0.36 | 0.66 |
| 0.39 | 0.48 | 0.64 | 0.77 | 0.21 | 0.53 |
| 0.33 | 0.99 | 0.55 | 0.28 | 0.12 | 0.37 |
| 0.45 | 0.46 | 0.42 | 0.54 | 5.45 | 0.36 |
| 0.23 | 0.5 | 0.85 | 0.61 | 0.22 | 0.58 |
| 0.27 | 0.49 | 0.43 | 0.89 | 0.16 | 0.09 |
| 0.16 | 0.43 | 0.32 | 0.51 | 0.4 | 1.03 |
| 0.24 | 0.33 | 0.45 | 0.34 | 0.32 | 0.42 |
| 0.29 | 0.3 | 0.53 | 0.34 | 0.3 | 0.43 |
| 0.42 | 0.46 | 0.54 | 0.3 | 0.24 | 0.32 |
| 0.18 | 2.96 | 0.44 | 0.48 | 0.21 | 0.25 |
| 0.22 | 0.14 | 0.77 | 0.67 | 0.16 | 0.41 |
| 0.35 | 0.72 | 0.57 | 0.63 | 0.12 | 0.81 |
| 0.39 | 0.45 | 0.81 | 0.61 | 0.35 | 0.79 |
| 0.42 | 0.36 | 0.7 | 0.75 |  | 0.4 |
| 0.23 | 0.23 | 0.48 | 0.32 |  | 0.28 |
| 0.33 | 1.42 | 0.39 | 0.51 |  | 0.29 |
| 0.28 | 0.61 | 0.59 | 0.67 |  | 0.44 |
| 0.29 | 0.28 |  | 0.88 |  | 0.35 |
| 0.39 | 0.54 |  | 0.79 |  | 0.25 |
| 0.28 | 0.38 |  | 0.49 |  | 0.31 |
| 0.28 | 0.2 |  | 0.48 |  | 0.46 |
| 0.33 | 1.26 |  | 1.95 |  | 0.31 |
| 0.48 | 0.37 |  | 0.56 |  | 0.35 |
| 0.25 | 0.41 |  | 0.62 |  | 0.51 |
| 0.13 |  |  | 0.68 |  | 0.52 |
| 0.21 |  |  |  |  | 0.29 |
| 0.25 |  |  |  |  | 0.19 |
| 0.26 |  |  |  |  | 0.51 |
| 0.37 |  |  |  |  | 0.26 |
| 0.21 |  |  |  |  | 0.33 |
| 0.21 |  |  |  |  | 1.38 |
| 0.31 |  |  |  |  | 0.36 |
| 0.31 |  |  |  |  | 0.5 |
| 0.51 |  |  |  |  | 0.72 |
| 0.22 |  |  |  |  | 0.39 |
| 0.35 |  |  |  |  | 0.34 |
| 0.18 |  |  |  |  | 1.99 |
| 0.29 |  |  |  |  |  |
| 0.26 |  |  |  |  |  |

**Figure 4f**

|  |  | ADP-ribosylation level | | |
| --- | --- | --- | --- | --- |
|  | Peptide, µM | Repeat 1 | Repeat 2 | Repeat 3 |
|  | 0 | 1 | 1 | 1 |
| F-tractin_opt_ | 1 | 1,245716 | 0,95033 | 1,419998 |
|  | 3 | 1,210256 | 1,296322 | 1,29383 |
|  | 10 | 0,846657 | 0,728925 | 1,078508 |
|  | 30 | 0,320904 | 0,220558 | 0,140877 |
|  | 100 | 0,032604 | 0,100767 | 0,002262 |
| F29A F-tractin_opt_ | 100 | 0,715886 | 1,144903 | 1,162086 |
| Lifeact | 1 | 0,978733 | 0,811616 | 1,268379 |
|  | 3 | 0,825802 | 0,671345 | 1,323177 |
|  | 10 | 0,671724 | 1,132312 | 0,970817 |
|  | 30 | 0,313245 | 0,438167 | 0,331448 |
|  | 100 | 0 | 0,032044 | 0,093063 |

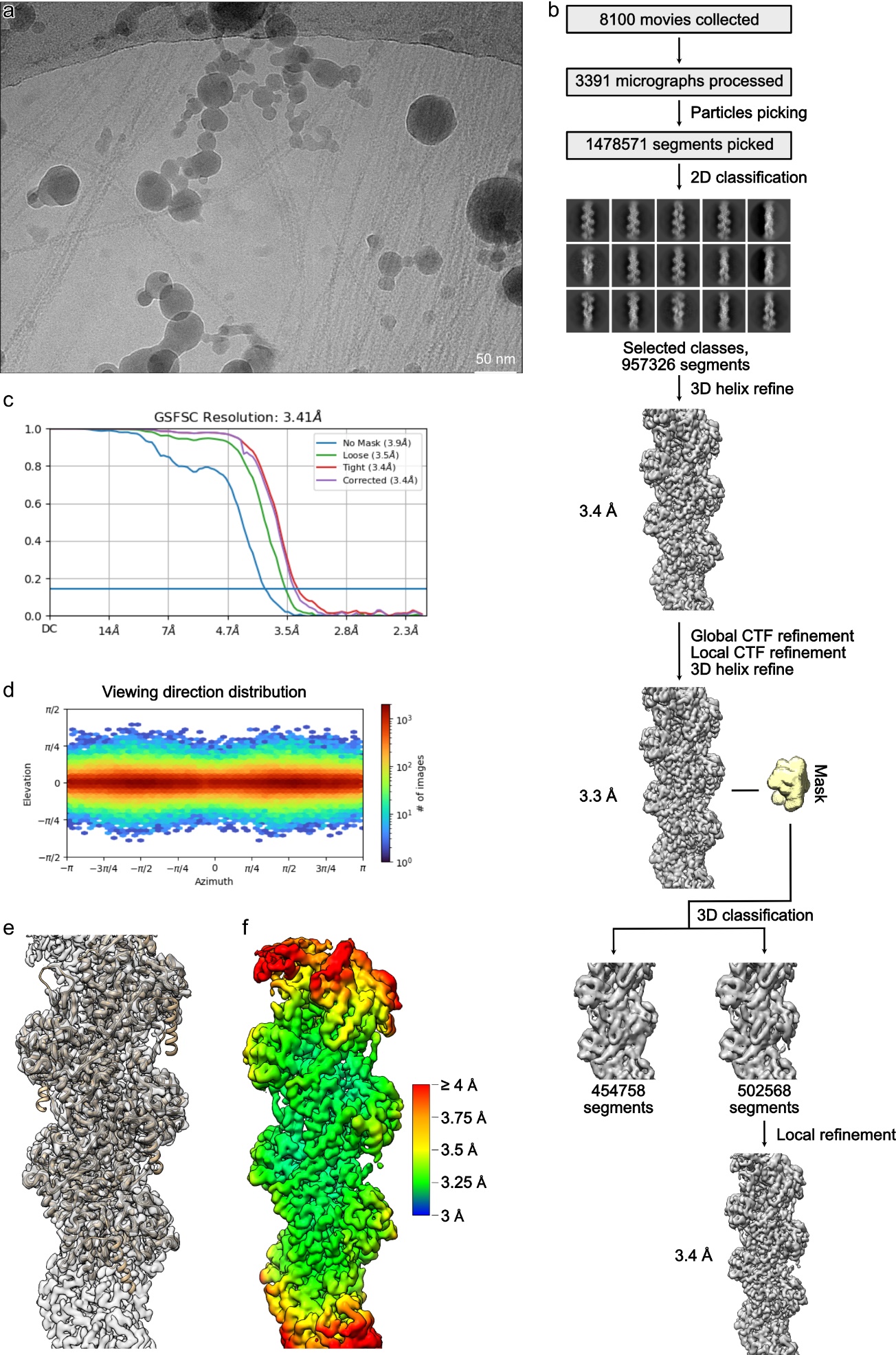

**Supplementary Fig. 1. Processing of the F-tractin-F-actin complex.** (a) An example of the 3391 analyzed cryo-EM micrographs.(b) Processing overview. (c) Fourier shell correlation curve of the final reconstruction. (d) Angular distribution. (e) Fit of the molecular model into the cryo-EM density. f Local resolution gradient of the reconstruction.

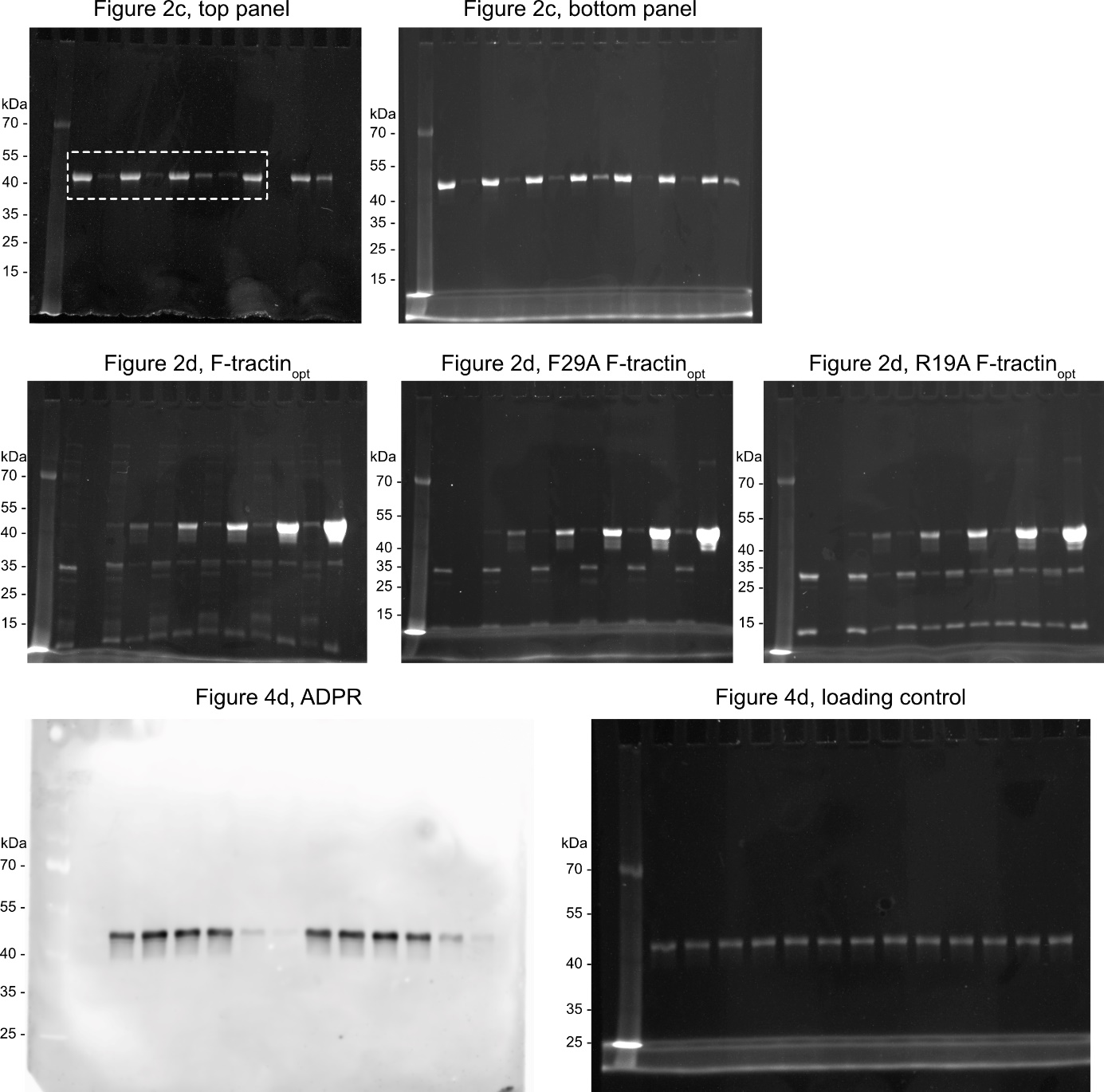

**Supplementary Fig. 2. Uncropped SDS-PAGE and western blots.**
